# Role of Chemokine and TNF signaling pathway in oral squamous cell carcinoma: A RNA deep sequencing analysis of oral buccal mucosa squamous carcinoma model of Chinese hamster

**DOI:** 10.1101/778399

**Authors:** Guoqiang Xu, Jianing Wei, Bing Huangfu, Jiping Gao, Xiaotang Wang, Lanfei Xiao, Ruijing Xuan, Zhaoyang Chen, Guohua Song

## Abstract

Oral cancer is one of the most common cancers in the world, meanwhile, differentially expressed genes are thought to regulate the development and progression of oral squamous cell carcinomas (OSCC). In this study we screened RNA transcripts from the oral buccal mucosa of healthy male Chinese hamster, divided into 3 groups: a control group with no disposal, a solvent control group coated with acetone solvent, and an experimental group coated with 0.5% DMBA acetone solution by high-throughput RNA sequencing. Tophat and Bowtie were used to align the high-quality reads into transcripts, DEseq was used to analysis the expression of differential gene. Then, the Gene Ontology (GO) enrichment analysis and Kyoto Encyclopedia of Genes and Genomes (KEGG) pathway enrichment analyses were conducted. The chemokine and TNF signaling pathway were differentially expression and the mRNA expression of *CXCL1*, *CXCL2*, *CXCL3*, *CCL7*, *MMP9*, monitored by qRT-PCR, increased remarkably in the cancer group and coincided with the result of RNA-Sequencing. Meanwhile, the *CXCL1*, *CXCL2*, *CXCL3*, and *CCL7* are significantly enriched in the chemokine signaling pathway, and *CXCL1*, *CXCL2*, *CXCL3*, and *MMP9* are significantly enriched in the tumor necrosis factor (TNF) signaling pathway. The differentially expression of the chemokine and TNF signaling pathway was a response to the invasion of the organism immune system due to oral buccal mucosa squamous carcinoma. All the findings provided novel insights for further molecular researches of oral cancer.

## Introduction

Oral cancer is one of the most frequent solid cancers worldwide, and oral squamous cell carcinoma (OSCC) constitutes around 90% of oral cancers (Siegel et al., 2014). It is highly invasive and metastatic at the advanced stage, and presents a substantial threat to human health. Meanwhile, there are 145000 deaths of OSCC in the world (1.8% of all cancer deaths in the world) annually; including 77% of the burden is in developing countries (Ferlay et al., 2015). Moreover, the incidence rate of OSCC is increasing, especially in younger people. Furthermore, OSCC has a very poor prognosis due to its invasive nature, the survival rate of patients with OSCC has not improved despite the improvements and innovations in diagnostic techniques and treatments (Shlok et al., 2010). However, clinical samples are pretty difficult to obtain and the less number of clinical samples meeting the experimental requirements is a major problem in oral cancer research, which seriously restricts the development of research on the mechanism of OSCC. Therefore, there is an urgent need for a better animal model of human OSCC lesions to help us better understand the pathogenesis of oral cancer.

The hamster cheek pouch is the most relevant known animal system that closely related to the human oral tumor like morphogenesis, phenotype markers and genetic alterations (Raimondi et al., 2005), meanwhile, the hamster buccal pouch mucosa is covered by a thin layer of keratinized stratified squamous epithelium that is similar in its thickness to the floor of the mouth and the ventral surface of the tongue in humans (Gimenez-Conti IB and TJ, 1993). Chinese hamster (Cricetulus griseus) also called striped-back hamster, is easy to operate by a single hand, has a buccal pouch on the each side of oral cavity. The establishment of animal models of Chinese hamster oral cancer can help us better study the development of oral cancer. In view of the above advantages, Chinese hamster becomes the ideal animal model of research on OSCC mechanism.

Recently, the studies by Shih-Han Lee shows that, in the process of cancer, even if the DNA itself has not changed, the inactivation or alteration of the tumor suppressor gene may also generate. Further studies have shown that alterations in cancer obtained during mRNA processing can essentially simulate DNA changes in somatic cells, and patients also have tumor suppressor gene inactivation, which indicate that cancer diagnosis of DNA alone may ignore other important molecular information that promotes disease progression (Lee et al., 2018). In common with other cancers, the occurrence and progression of OSCC is a multistep process with the accumulation of genetic and epigenetic changes (Kang and Park, 2001). Evidence from various molecular and genetic studies suggests the association between squamous cell carcinoma initiation and development and the accumulation of genetic alterations at both the DNA and RNA levels (Gibb et al., 2010). Recent studies have indicated that the changes of mRNA expression levels in OSCC are associated with tumor development, maintenance, and progression. In addition, mRNAs could be potentially used as biomarkers of OSCC or other oral cancers (Lodi et al., 2010).

Compared to traditional techniques, next generation sequencing (NGS) offers greatly improved dynamic ranges and specificity for transcriptome analyses, while sample throughput is continuously increasing and costs are being reduced, and gives a far more precise measurement of transcript expression levels and a far more sophisticated characterization of transcript isoforms, even a far more reliable characterization of allele specific expression patterns (Christopher et al., 2010). Although there have been published reports on the successful construction of genome-wide mRNA expression profiles in other types of cancer, highlighting the power and capability of high-throughput sequencing techniques, there have been relatively few relating to OSCC (Li et al., 2011; Zhu et al., 2011). Herein, high-throughput RNA sequencing on tumor samples and their matched normal samples combined with qRT-PCR analysis were used to investigate the mechanisms of the occurrence and development of OSCC. We used the information to explore and predict the molecular functions and biological pathways of the differentially expressed mRNAs. Our results have provided a basis for identifying further molecular markers for the diagnosis and treatment of OSCC.

## Materials and Methods

### Establishment of animal model

Sixty male Chinese hamsters (Cricetulus griseus, aged 8-10 weeks, 21-25g b.w.) were provided by the Experimental Animal Center of Shanxi Medical University (Taiyuan, China SCXK [Jin] 2015-0001), housed in standard hamster cages, maintained in a temperature-controlled environment with a 12 h light/dark cycle and fed a standard hamster diet and water ad libitum(SYXK [Jin] 2015-0001). After one-week acclimation, these hamsters were divided randomly into three groups: treatment group (24 hamsters), which was coated the buccal pouch with 0.5% 7,12-dimethylbenz[a]anthracene (DMBA) acetone solution; solvent control group (12 hamsters), which was only coated with acetone solution; control group (24 hamsters), which was subjected to no disposal. The doses were chosen on the basis of the previous studies, and to observe the entire carcinogenesis of oral cancer, animals were treated with DMBA three times a week for 15 weeks (Rajasekar et al., 2016; Ramu et al., 2017). All experimental procedures were conducted and performed during the light cycle in accordance with the Animal Care and Use Committee regulations of Shanxi Medical University.

### Histopathological analysis

The tumor tissues were fixed in 4% paraformaldehyde for about 48 hours, transferred to 70% ethanol, then processed in a graded series of ethanol solutions, embedded in paraffin and cut into 4μm thick sections. The sections were stained with hematoxylin and eosin (H & E) for histological examination. The histopathological specimens were observed under light microscope by oral pathologist experts. The pouch pathological changes were determined according to the 12 grade record in the WHO standard (None, 1978).

### Ultrastructure of oral buccal pouch mucosa observation

The oral buccal mucosa was extracted, washed with physiological saline, cut into 1-mm^3^ pieces, fixed in 2.5% glutaraldehyde for 2h at 4°C dehydrated through a graded series of ethanol, embedded in epoxy resin, trimmed, sectioned, and observed and photographed under a JEM-1011 transmission electron microscope(Tokyo Japan).

### Enzyme-linked immunosorbent assay (ELISA)

In order to investigate the serum levels of TNF-α, AFP, and SCC-Ag in serum of Chinese hamster with and without OSCC, we collected about 1 ml blood from each animal prior to surgery. Blood samples were allowed to clot for 30 min at room temperature (RT) and serum was obtained by centrifugation at 1300 g for 10 min at 4°C. Serum samples were then aliquoted and stored at − 80°C.

Serum samples were thawed on ice, the levels of TNF- α (Cat. MM-160001, MEIMIAN, China), AFP(Cat. JL-F46753, JiangLai, China), and SCC-Ag(Cat. JL-F46768, JiangLai, China) were detectable with the commercially available enzyme-linked immunosorbent assay (ELISA) kit according to manufacturer’s instructions.

### RNA Sample Preparation

All Chinese hamsters were killed by cervical dislocation on the week 15, six pairs of OSCC and normal buccal pouch samples of Chinese hamster, which had been identified by pathologists, were immediately isolated, frozen in liquid nitrogen and stored at −80 °C for RNA extraction and gene expression research. Total RNA was extracted from 6 pairs of OSCC and normal buccal pouch samples using TRIzol (Invitrogen, USA) according to the protocol provided by the manufacturer. The purity and integrity of total RNA were monitored by NanoPhotometer® spectrophotometer (Implen, GER) and the Agilent 2100 Bioanalyzer (Agilent, USA), respectively.

### RNA Library Construction and Deep Sequencing

Briefly, mRNA was isolated from total RNA using the FastTrack MAG mRNA Isolation Kits according to the manufacturer’s instructions (Invitrogen, USA). Before doing any further steps, Fragmentation Buffer (Agilent, USA) was applied to perform fragmentation on qualified mRNA coming from buccal pouch samples of Chinese hamster. Then, the mRNA was converted into double-stranded cDNA by reverse transcription. Following end repair and A-tailing, paired-end sequencing adaptors complementary to sequencing primers were ligated to the ends of the cDNA fragments. The ligated products were purified on 2% agarose gels, qualified fragments were selected for downstream enrichment by PCR. Each sample was subjected to sequencing from both ends by the Illumina Hiseq 2500 sequencing technology.

### Data Filtering

Raw image data received by sequencing were transformed by base calling (CASAVA) into sequence data (termed Raw Reads). Raw Reads were filtered through data processing to get Clean Reads. We removed artificial reads, adapter contamination reads. High-quality reads with a length of >50 bp were reserved.

### RNA-seq Reads Alignment

In our experiment, Tophat v2.0.12 was used to align the reads into transcripts based on the Cricetulus griseus reference genome in the UCSC (University of California Santa Cruz) genome database, and Bowtie2 was used to comparison analysis.

### Differential Expression Analysis

DEseq was used to analysis the expression of differential gene, compare the expression of the same gene in normal buccal pouch and buccal pouch of OSCC in Chinese hamsters, and select genes with |log_2_Ratio| ≥ 1 and Corrected *P*-value < 0.05 as differentially expressed gene to obtain up-regulated and down-regulated genes.

### Gene Ontology Functional Analysis

For gene expression profiling analysis, Gene Ontology (GO) enrichment analyses of functional significance were performed by hypergeometric testing to map all differentially expressed genes to GO terms. The calculated *P*-value is corrected and FDR < 0.05 was used as the threshold to judge the differential gene expression of significantly enriched GO terms. The functional classification of the two groups of genes on the GO (Gene Ontology) was compared and the top ten of the three main ontology of GO term were selected for GO term enrichment analysis.

### KEGG Pathway Analysis

The KEGG database was used to assign the assembled sequences (http://www.genome.jp/kegg), which can facilitate understanding of the biological functions of genes and identify significantly enriched metabolic pathways or signal transduction pathways in differentially expressed genes. The method of KEGG pathway analysis was the same as the GO analysis, and the significant enrichment pathway was analyzed by KAAS. FDR was used to correct the related parameters, pathways with FDR ≤ 0.05 were considered significantly enriched pathway.

### Quantitative Real Time PCR

Quantitative real-time PCR (qRT-PCR), the traditional quantification method on gene expression, was adopted to further confirm the findings from the RNA-seq analysis. On the basis of the results, we filtered five significant differentially expressed genes involved in chemokine signaling pathway and TNF signaling pathway. According to Chinese hamster *CXCL1* sequence (Accession No: NM_001244044.1), *CXCL2* sequence (Accession No: XM_003510006.2), *CXCL3* sequence (Accession No: NM_001244139.1), *CCL7* sequence (Accession No: XM_003495790.2), *MMP9* sequence (Accession No: XM_007641300.1) and *β-actin* sequence (Accession No: NM_001244575.1) in GenBank, primers were designed with Primer Premier 5 software, evaluated by Oligo 6 software and subjected to Blast specificity tests at NCBI. β-actin was used as the endogenous control and all the primers were synthesized in Huada Gene. Primer sequence information is as Table 1.

**Table1.**
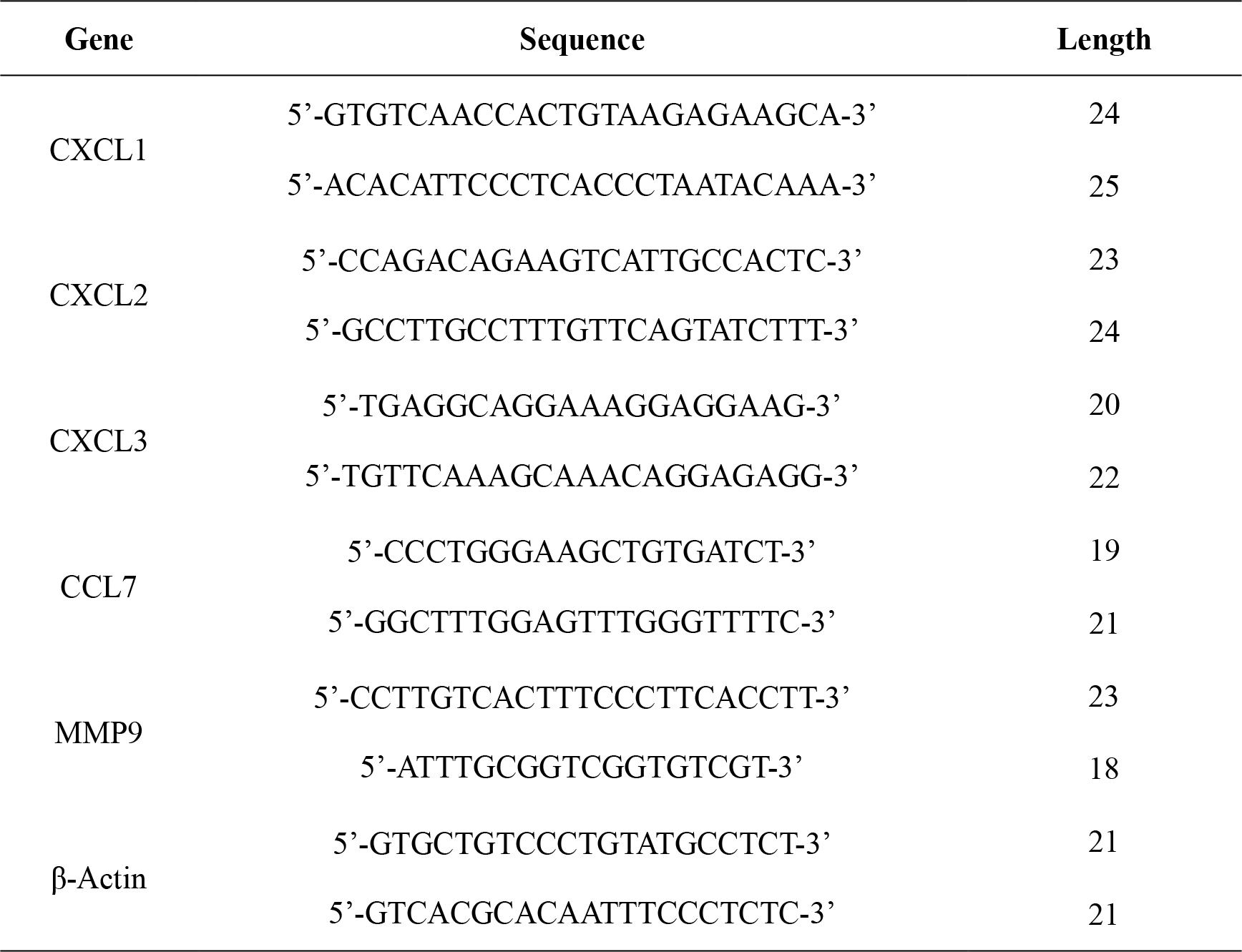
Primers sequence information.

The qRT-PCR analysis was conducted in a total volume of 20μL containing 10μl 2 × SYBR® Premix Ex TaqII (Tli RNaseH Plus) (Takara Bio Inc., China), combined with sense and antisense primers (0.8 μl, final concentration 0.4 μM), ddH_2_O 6.4 μl, and 2 μl diluted cDNA in Optical 8-Tube Strip using the Applied Biosystems 7300 Real-Time PCR Instrument (ABI, USA). The conditions for real-time PCR were as follows: after initial denaturation at 95°C for 30 s, 40 PCR cycles were started with thermo cycling conditions at 95°C for 5 s, 60°C for 30 s, and then a melting curve analysis was performed to verify the specificity of the PCR product. Every sample was analyzed in triplicate. System software and the 2^−ΔΔCt^ method were applied to quantitative calculation.

### Data Analysis

Statistical analysis was performed using SPSS v17.0 software and Student’s t-test was used to analyze the OSCC and normal tissue samples. All values in the experiment were expressed as mean ± SEM (standard error of the mean) and values of *P* < 0.05, were expressed statistically significant.

## Result

### The effect of DMBA-induced oral carcinogenesis

To identify cancer tissues of oral buccal pouch mucosa cancer model of Chinese hamster, we examined the histological changes of buccal pouch mucosa by HE stain (Figure 1). It’s important to note that in our research no significant differences were observed in solvent control group compared with the control group after 15 weeks acetone solution treatment. Meanwhile, when dissecting, we found that buccal pouch of the cancer group became embrittled and shrinking volume. Furthermore, compared with the control group, we observed the cancer group showed atypical nuclear division with initial keratinization and enlarged nucleoli, cells broke through the basement membrane, infiltrating the lamina propria and connective tissue, and tumor islands emerged, was highly differentiated squamous cell and invasive cancer. All of these provided the support for sequence analysis.

**Figure 1.**
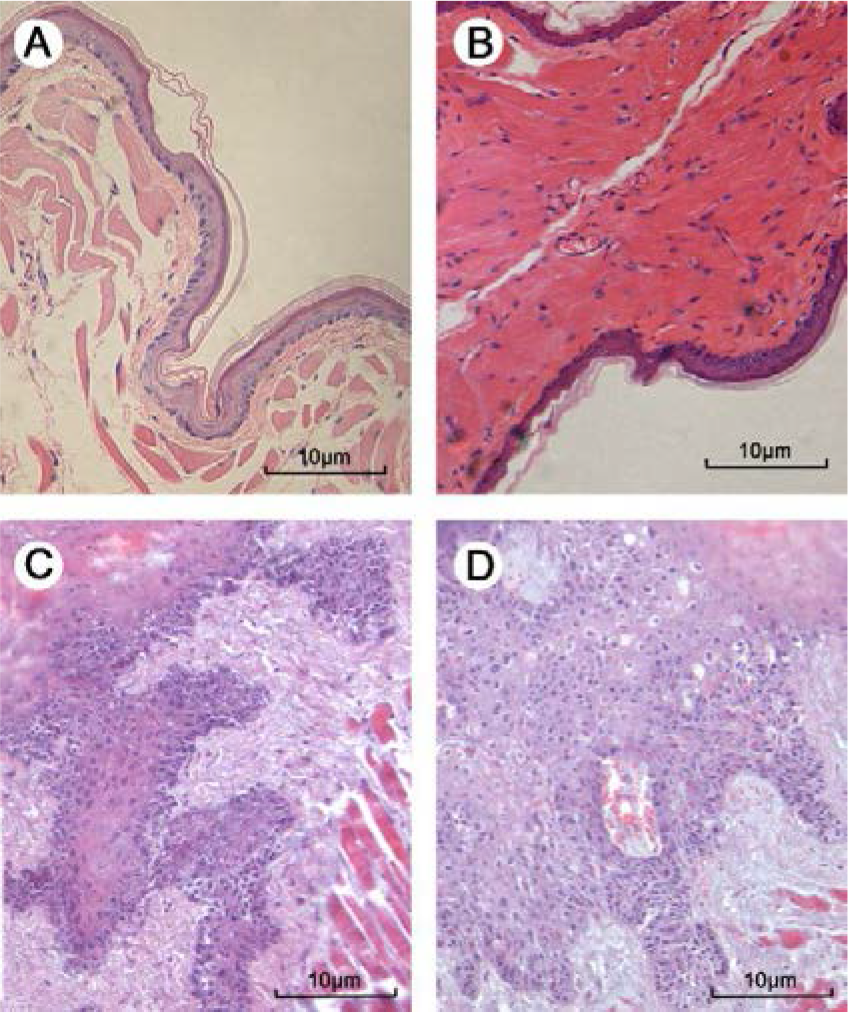
Pathological analysis of oral buccal mucosa between control, solvent control and cancer group. A: control group (×200). B: solvent control group (×200). C & D: cancer group (×200). A & B showed that there is a thin layer of connective tissue between the epithelium and the muscular layer, and the basal cells are arranged in a neatly arranged shape. Meanwhile, in C & D epithelial cells and nucleus show significant pleomorphism, cells break through the basement membrane, infiltrate the lamina propria and connective tissue.

### The change of oral buccal pouch mucosa ultrastructure

In order to further understand the alterations in the buccal pouch of the cancer group, we used transmission electron microscope to detect them (Figure 2). Compared with the control group, in the cancer group, the shape of the cells is irregular, the nucleolus is concentrated, the nucleus is jagged, the nuclear membrane is invaginated, and the desmosome is reduced or even disappeared.

**Figure 2.**
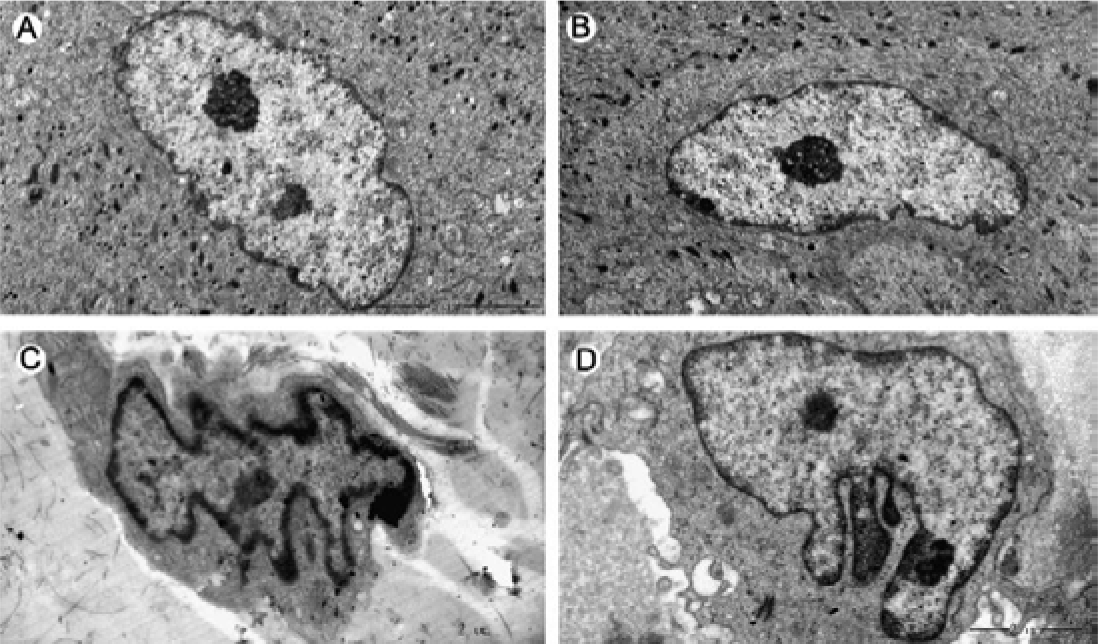
Ultrastructure analysis of oral buccal pouch mucosa between control, solvent control and cancer group. A: control group (×8000). B: solvent control group (×8000). C & D: cancer group (×12000). A & B showed that the shape of the cells is regular and closely arranged, the shape of the nucleus is regular, and the morphology of each organelle is normal and the desmosome is abundant. At the same time, the C & D showed that the shape of the cells is irregular, the nuclear condensation into jagged, the nuclear membrane is invaginated, and the desmosome is reduced or even disappears.

### The levels of the TNF- α, AFP, and SCC-Ag

Measures of TNF-α, AFP, and SCC-Ag in serum of OSCC Chinese hamsters were shown in Figure 3. The level of serum TNF-α in Chinese hamsters with OSCC was significantly lower than healthy control group (P<0.01). While, the level of serum SCC-Ag was higher than healthy control group (P<0.05). Among them, AFP values were not significantly different between control and OSCC.

**Figure 3.**
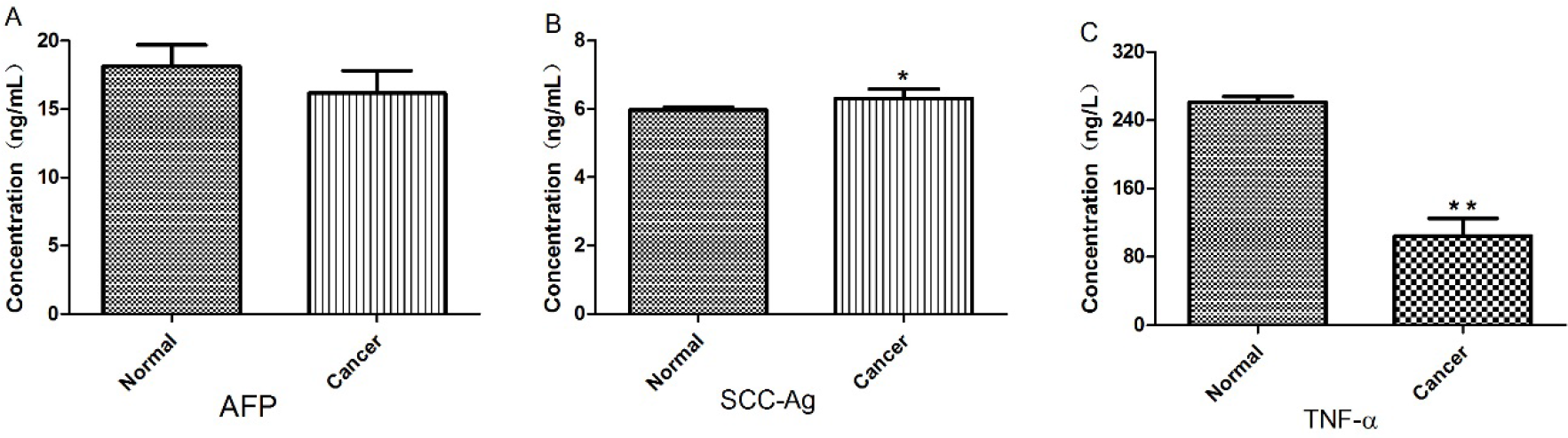
The levels of the AFP, SCC-Ag, and TNF-α in oral buccal pouch mucosa between control and cancer group. TNF-α in Chinese hamsters with OSCC was significantly lower than healthy control group (P<0.01). While, the level of serum SCC-Ag was higher than healthy control group (P<0.05). Among them, AFP values were not significantly different between control and OSCC.

### Overview of Sequencing Data from RNA-seq Analysis

In both the cancer and normal groups, about (32 ~ 44)× 10^6^ reads (73 ~ 92% of the total raw reads) were aligned to *Cricetulus griseus* reference genome sequence among samples, with an average of 38×10^6^ reads per sample. The unique alignment sequence of the two groups accounts for 70 ~ 90% of the reference genome. The distribution of the unique alignment sequences in the reference genome was shown in Figure 4.

**Figure 4.**
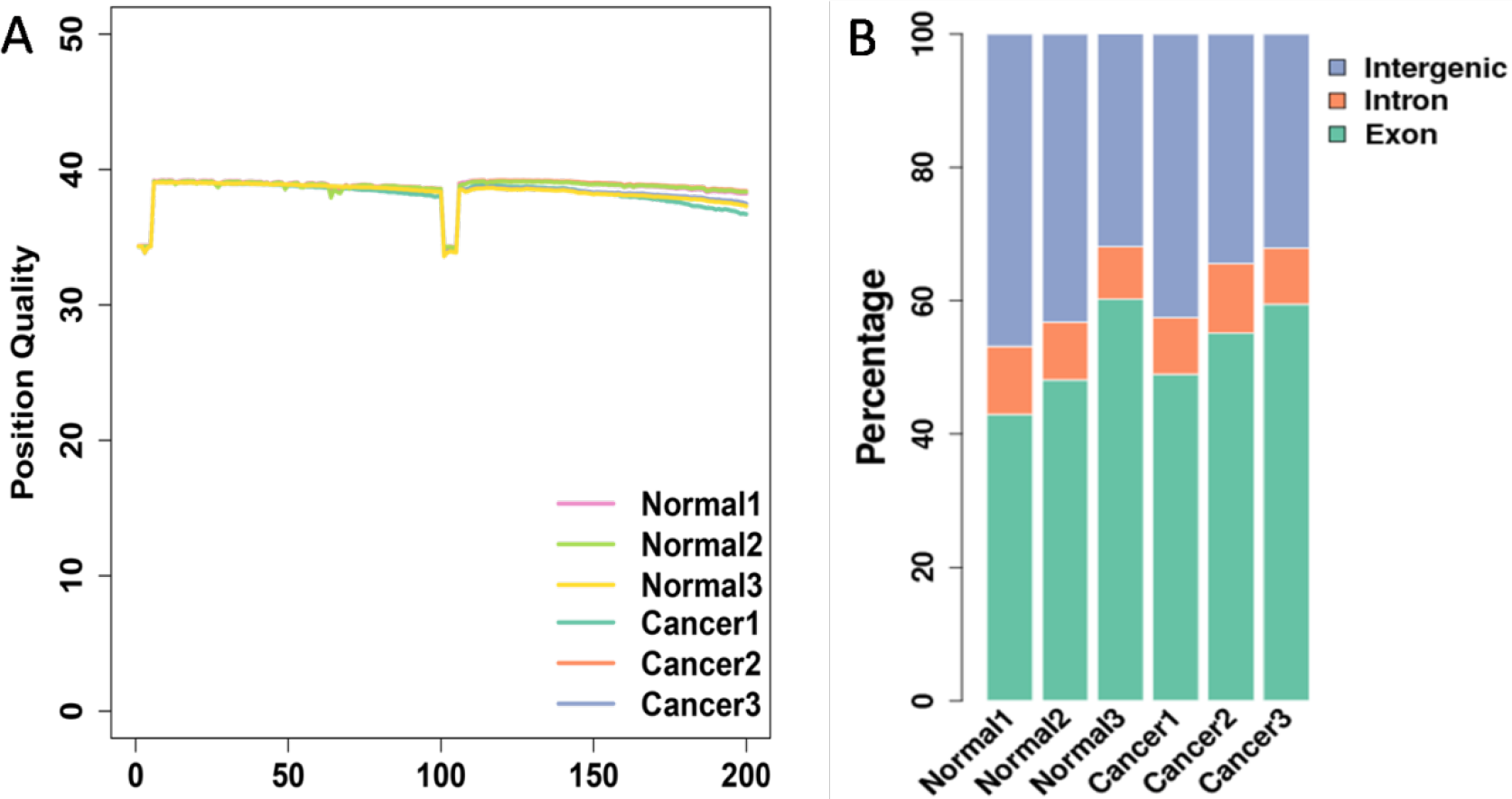
Sequence quality and unique alignment sequence distribution of cancer and normal groups. **A:** Sequencing quality distribution of the samples. **B:** The distribution of the unique alignment sequence in each region of genes in the reference genome.

### Analysis of Differentially Expressed Genes

Analysis of the data indicated that there were 194 significantly differentially expressed genes in the treated samples, of which 66 (34.02%) were down-regulated and 128 (65.97%) were up-regulated (Figure 5). The top five up-regulated and down-regulated genes with the significant changes in expression were shown in the Figure 6A, and the top thirty of the altered genes was listed in Table 2. The hierarchical cluster analysis of differentially expressed genes in Cancer and Normal groups revealed that multiple gene modules were formed by many genes with similar expression trends, and these genes may be involved in the occurrence, metastasis, and invasion of oral squamous cell carcinoma (Figure 6).

**Table2.** Top thirty genes altered in cancer group.

| Gene | Log<sub>2</sub> Ratio | Corrected *P*-<br>value* | Gene | Log<sub>2</sub> Ratio | Corrected *P*-<br>value* |
| --- | --- | --- | --- | --- | --- |
| CXCL2 | 26.69 | 5.54E-15 | CCL7 | 5.88 | 2.13E-05 |
| MMP13 | 11.69 | 1.18E-14 | IL1B | 11.53 | 2.13E-05 |
| CCL3 | 26.24 | 4.13E-14 | PTX3 | 7.31 | 2.38E-05 |
| LCN1 | 25.19 | 1.27E-10 | CAIV | 5.85 | 2.38E-05 |
| MMP9 | 7.42 | 4.35E-09 | CXCL7 | 22.94 | 4.54E-05 |
| CXCL3 | 6.52 | 1.42E-07 | OLR1 | 22.97 | 4.61E-05 |
| STRA6 | 7.15 | 1.68E-07 | RN45s | -8.19 | 5.53E-05 |
| LOC100754872 | 23.63 | 6.98E-07 | THBS1 | 4.72 | 6.72E-05 |
| Serpine1 | 6.05 | 8.31E-07 | IL-36 γ | 6.67 | 9.12E-05 |
| TNN | 8.20 | 1.23E-06 | CXCL1 | 5.77 | 0.0001 |
| CTHRC1 | 8.46 | 1.35E-06 | LOC100766767 | -7.59 | 0.0001 |
| GPx6 | -24.40 | 4.67E-06 | MS4A7 | 5.99 | 0.0001 |
| CTRP6 | 5.52 | 6.88E-06 | SFRP2 | 5.02 | 0.0001 |
| LOC100764819 | -8.33 | 1.40E-05 | POSTN | 4.81 | 0.000115 |
| SLN | -6.62 | 1.49E-05 | MMP12 | 5.23 | 0.000121 |

**Figure 5.**
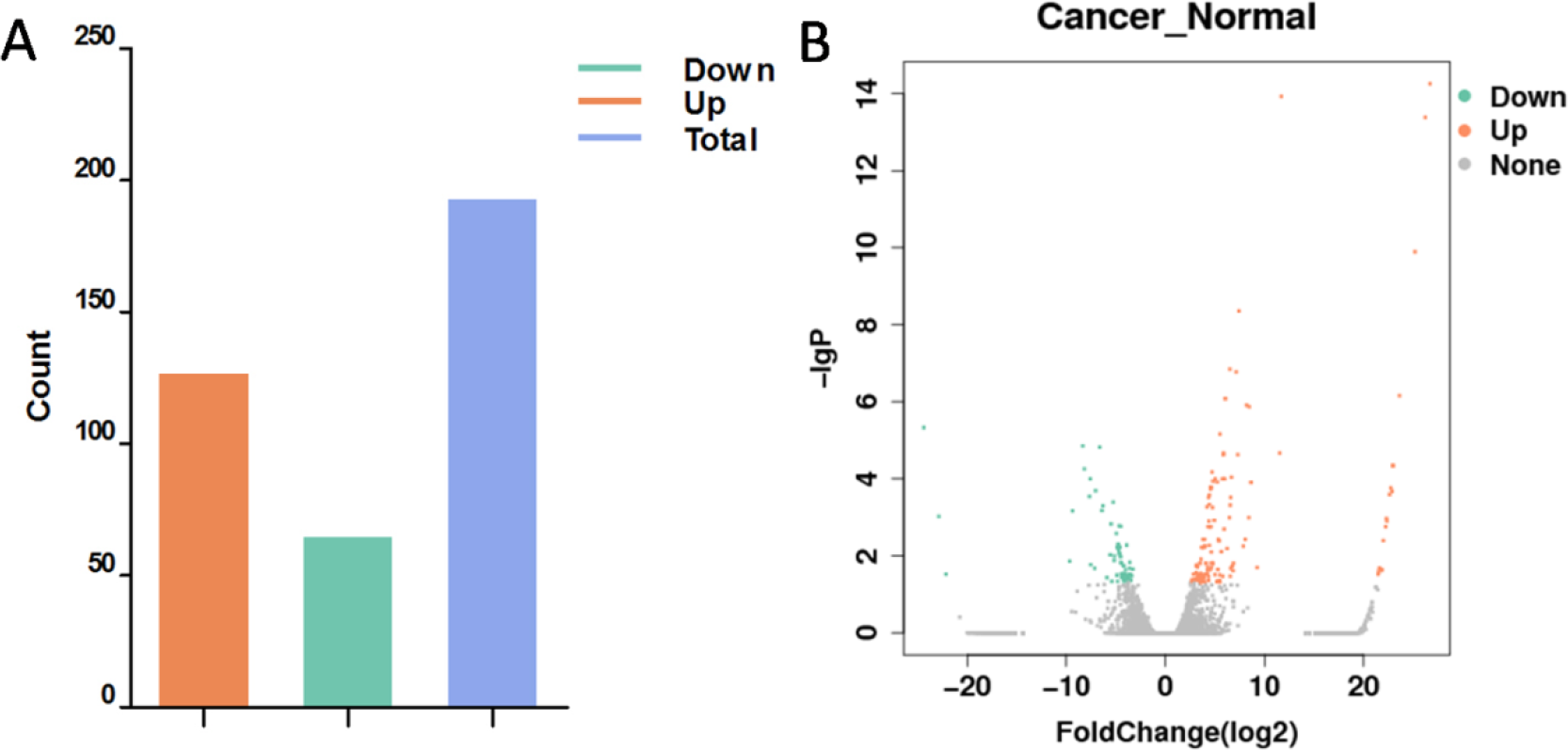
Histograms and volcano plot differentially expressed gene in cancer and normal groups. A: Histogram, Y-axis is the amount of differentially expressed genes, X-axis is the classification of differentially expressed genes, red indicates up-regulated genes, green indicates down-regulated genes, and blue indicates total differentially expressed genes. B: volcano plot, Y-axis is −log10 (P-value) of differentially expressed genes, X axis is the log2 FoldChange of the differentially expressed genes, red indicates up-regulated genes and green indicates down-regulated genes. There were 66 (34.02%) were down-regulated and 128 (65.97%) were up-regulated.

**Figure 6.**
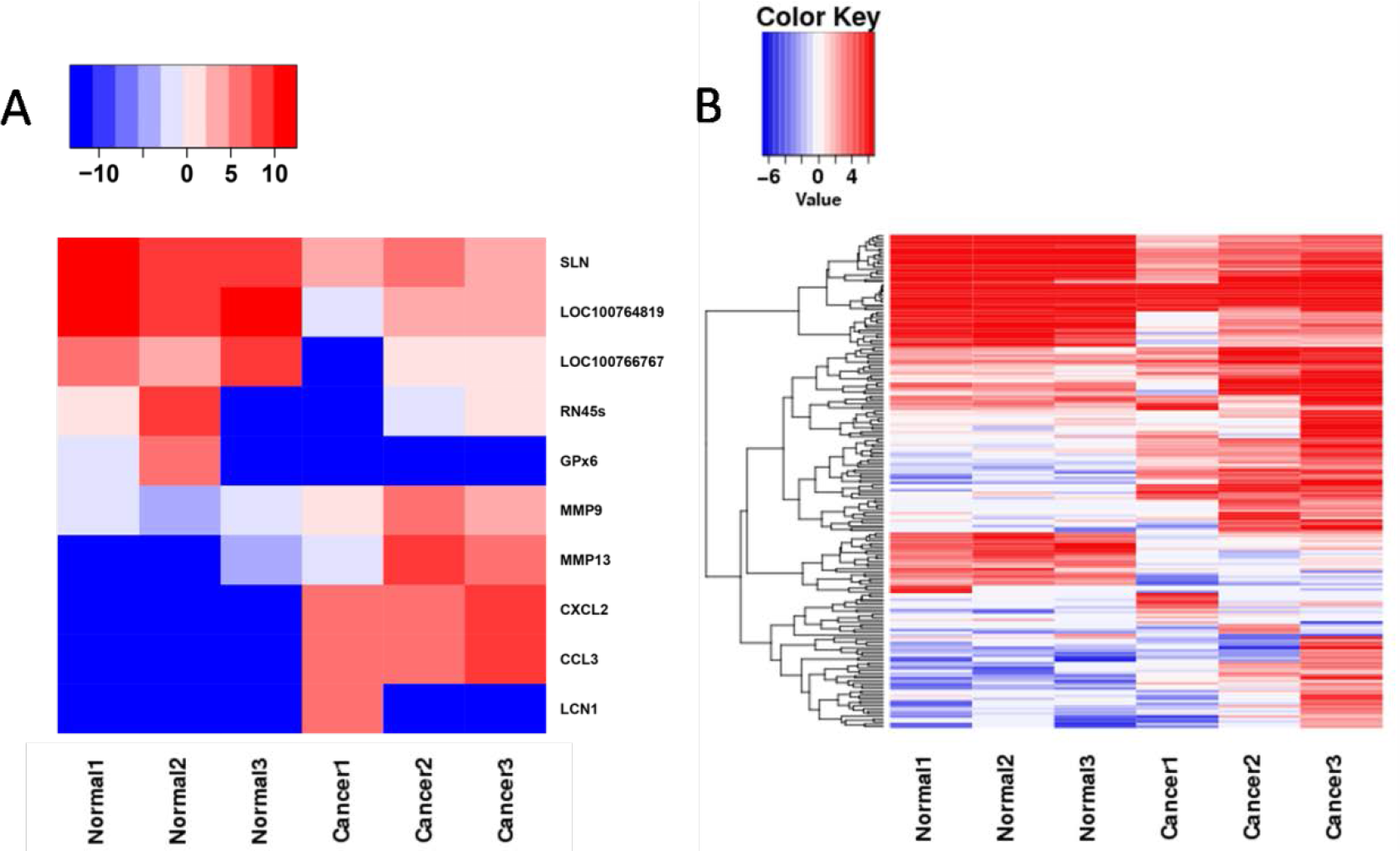
Hierarchical clustering of differentially expressed genes in oral squamous cell carcinoma and normal tissues. **A:** Hierarchical clustering graph, each rectangle represents the expression value of a certain gene (row) in a certain sample (column), and the gene expression changes from blue (low expression) to red (high expression). **B:** The top five up-regulated and down-regulated genes.

### Differential Expression GO Analysis

In the GO (Gene Ontology) statistics of differentially expressed genes, 184 genes were classified according to the GO classification method, and the amount of up-regulated and down-regulated differentially expressed genes in each subclass was calculated (Figure 7). The GO enrichment analysis of differentially expressed genes in cancer samples identified 120 biological processes, 18 cellular components, and 2 molecular functions that are closely related to OSCC and the top ten of them are listed in the Table 3.

**Table3.**
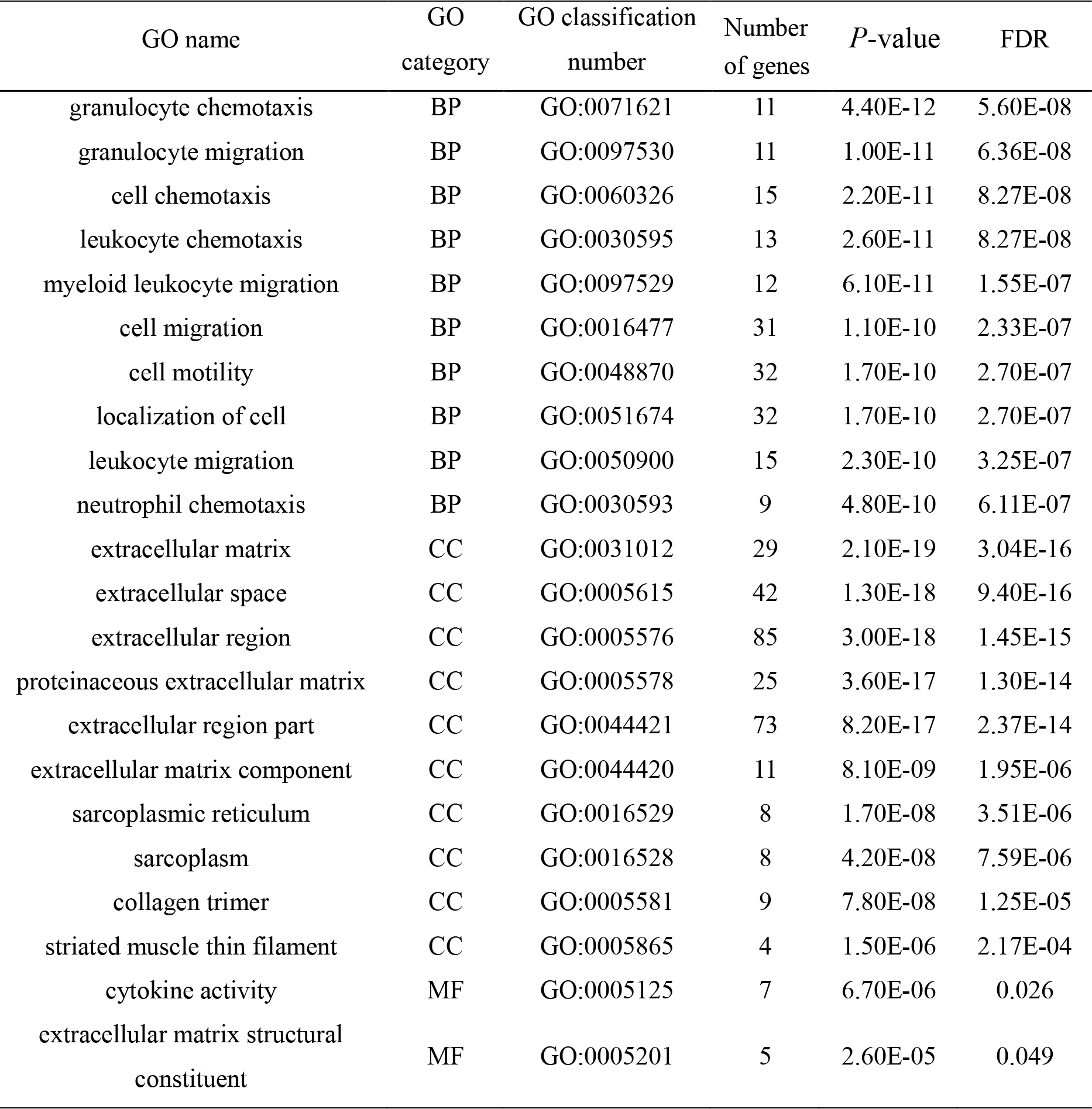
GO enrichment analysis of differentially expressed genes in cancer and normal groups.(FDR<0.05) (BP: Biological Process; CC: Cellular Component; MF: Molecular Function)

| GO name | GO category | GO classification number | Number of genes | P-value | FDR |
| --- | --- | --- | --- | --- | --- |
| granulocyte chemotaxis | BP | GO:0071621 | 11 | 4.40E-12 | 5.60E-08 |
| granulocyte migration | BP | GO:0097530 | 11 | 1.00E-11 | 6.36E-08 |
| cell chemotaxis | BP | GO:0060326 | 15 | 2.20E-11 | 8.27E-08 |
| leukocyte chemotaxis | BP | GO:0030595 | 13 | 2.60E-11 | 8.27E-08 |
| myeloid leukocyte migration | BP | GO:0097529 | 12 | 6.10E-11 | 1.55E-07 |
| cell migration | BP | GO:0016477 | 31 | 1.10E-10 | 2.33E-07 |
| cell motility | BP | GO:0048870 | 32 | 1.70E-10 | 2.70E-07 |
| localization of cell | BP | GO:0051674 | 32 | 1.70E-10 | 2.70E-07 |
| leukocyte migration | BP | GO:0050900 | 15 | 2.30E-10 | 3.25E-07 |
| neutrophil chemotaxis | BP | GO:0030593 | 9 | 4.80E-10 | 6.11E-07 |
| extracellular matrix | CC | GO:0031012 | 29 | 2.10E-19 | 3.04E-16 |
| extracellular space | CC | GO:0005615 | 42 | 1.30E-18 | 9.40E-16 |
| extracellular region | CC | GO:0005576 | 85 | 3.00E-18 | 1.45E-15 |
| proteinaceous extracellular matrix | CC | GO:0005578 | 25 | 3.60E-17 | 1.30E-14 |
| extracellular region part | CC | GO:0044421 | 73 | 8.20E-17 | 2.37E-14 |
| extracellular matrix component | CC | GO:0044420 | 11 | 8.10E-09 | 1.95E-06 |
| sarcoplasmic reticulum | CC | GO:0016529 | 8 | 1.70E-08 | 3.51E-06 |
| sarcoplasm | CC | GO:0016528 | 8 | 4.20E-08 | 7.59E-06 |
| collagen trimer | CC | GO:0005581 | 9 | 7.80E-08 | 1.25E-05 |
| striated muscle thin filament | CC | GO:0005865 | 4 | 1.50E-06 | 2.17E-04 |
| cytokine activity | MF | GO:0005125 | 7 | 6.70E-06 | 0.026 |
| extracellular matrix structural constituent | MF | GO:0005201 | 5 | 2.60E-05 | 0.049 |

**Figure 7.**
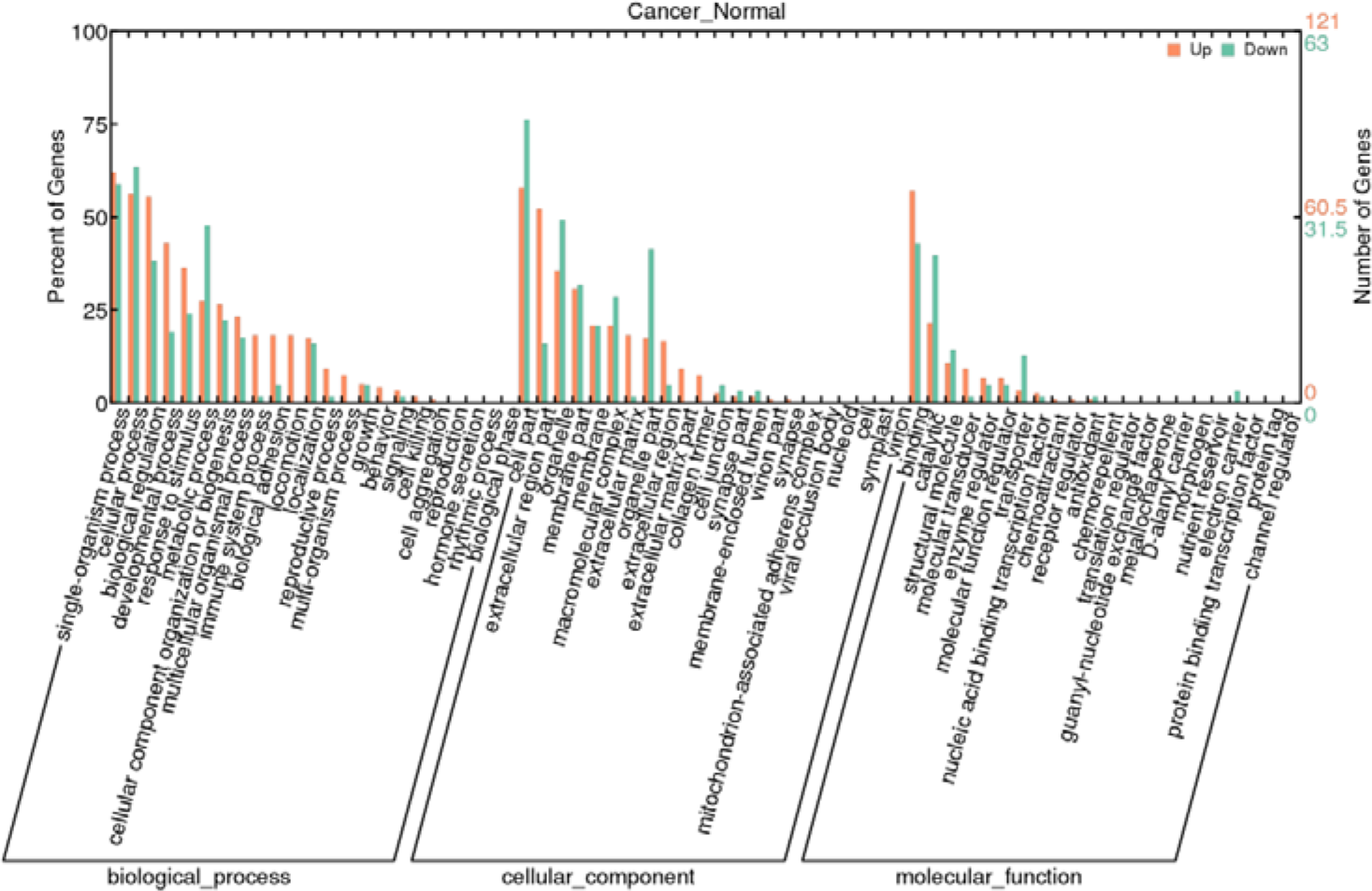
GO classification statistics histogram of differentially expressed genes between cancer and normal groups. Red indicates up-regulated genes, green indicates down-regulated genes, and all differentially-expressed genes are classified into 67 different subclasses (third-level entry) according to GO classification.

### KEGG Pathway Enrichment Analysis

Based on the analysis of KEGG pathway enrichment, we found that differentially expressed genes in the cancer group were mainly enriched in 8 signaling pathways including tumor necrosis factor pathway (TNF), chemokine pathway, cytokine interaction pathway, extracellular matrix receptor interaction pathway, protein digestion pathway and absorption pathway; and the number of enriched gene in the cytokine interaction pathway and chemokine pathway is the highest, suggesting that these pathways may play an crucial role in tumorigenesis, invasion and metastasis (Table 4).

**Table4.** Significant enrichment pathway of differentially expressed genes (Q-value<0.05)

| KEGG signal pathway | Map | Count | Q-value | Genes |
| --- | --- | --- | --- | --- |
| Malaria | map05144 | 5 | 0.002 | CSF3;IL1B;THBS2;SELP;CCL2 |
| ECM-receptor interaction | map04512 | 6 | 0.005 | LOC103164586;TNC;<br>THBS2;TNN;SPP1;COL1A1 |
| Salmonella infection | map05132 | 6 | 0.005 | CXCL1;CXCL3;CXCL2;IL1B;<br>CCL4; CCL3 |
| Chemokine signaling pathway | map04062 | 8 | 0.005 | CCL7;CXCL1;CXCL3;CXCL2;<br>CCL4;CCL3;CCL2;CXCL7; |
| Protein digestion and absorption | map04974 | 6 | 0.005 | LOC10316458; OL18A;COL7A;<br>COL12A;COL1A1; SLC1A1; |
| Cytokine-cytokine receptor interaction | map04060 | 8 | 0.006 | CSF3R;CCL7;CSF3;IL1B;<br>CCL4;CCL3;CCL2;CXCL7 |
| TNF signaling pathway | map04668 | 6 | 0.008 | CXCL1;CXCL3;CXCL2;<br>MMP9;IL1B;CCL2; |
| Toll-like receptor signaling pathway | map04620 | 5 | 0.047 | IL1B;CCL4;CCL3;SPP1;CTSK; |

We screened five differentially expressed genes related to OSCC, the Table 5 clearly reflects that CXCL1, CXCL2, CXCL3, CCL7, and MMP9 are enriched in multiple GO terms and KEGG pathways. As is depicted in the Figure 8(Kanehisa et al., 2017), CXCL1, CXCL2, CXCL3, and CCL7 are significantly enriched in the chemokine signaling pathway (Figure 8A), whereas CXCL1, CXCL2, CXCL3, and MMP9 are significantly enriched in the tumor necrosis factor signaling pathway (Figure 8B), suggesting that CXCL1, CXCL2, CXCL3, CCL7, and MMP9 may participate in the development of OSCC by molecular dialogues and interactions in metabolic pathways.

**Table5.** GO and KEGG enrichment pathways of 5 screened genes

| Gene | *P*-value | GO-BP | TOP 3 GO-BP<br>enrichment pathway | KEGG enrichment pathway |
| --- | --- | --- | --- | --- |
| CXCL1 | 0.0001 | 13 | cell chemotaxis | Salmonella infection |
|  |  |  | cell migration | Chemokine signaling pathway |
|  |  |  | cell motility | TNF signaling pathway |
| CXCL2 | 5.54E-15 | 41 | granulocyte chemotaxis | Salmonella infection |
|  |  |  | granulocyte migration | Chemokine signaling pathway |
|  |  |  | cell chemotaxis | TNF signaling pathway |
| CXCL3 | 1.42E-07 | 33 | granulocyte chemotaxis | Salmonella infection |
|  |  |  | granulocyte migration | Chemokine signaling pathway |
|  |  |  | cell chemotaxis | TNF signaling pathway |
| CCL7 | 2.13E-05 | 101 | granulocyte chemotaxis | Chemokine signaling pathway |
|  |  |  | granulocyte migration | Cytokine-cytokine receptor interaction |
|  |  |  | cell chemotaxis |  |
| MMP9 | 4.35E-09 | 60 | cell migration |  |
|  |  |  | cell motility | TNF signaling pathway |
|  |  |  | localization of cell |  |

**Figure 8.**
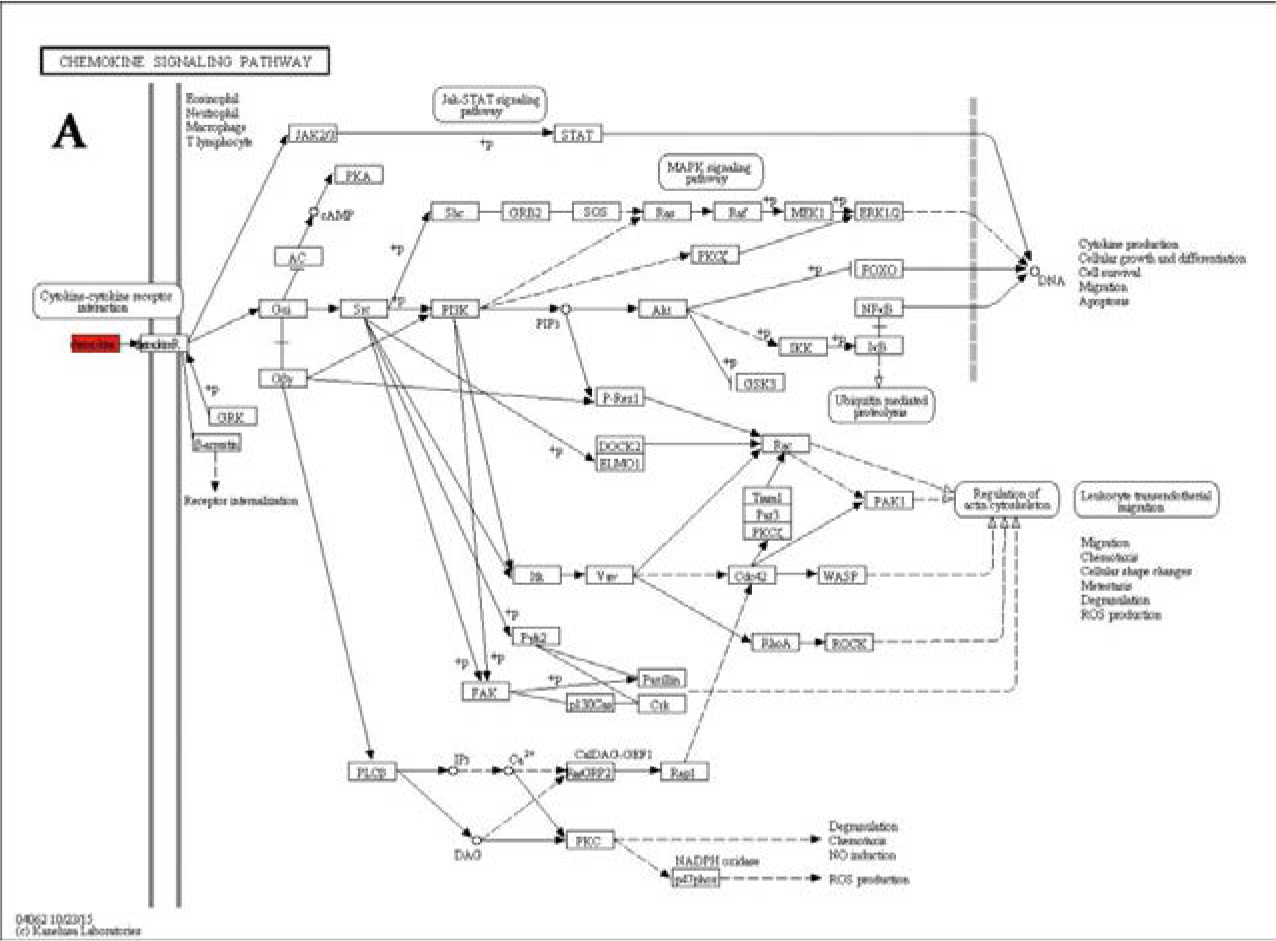
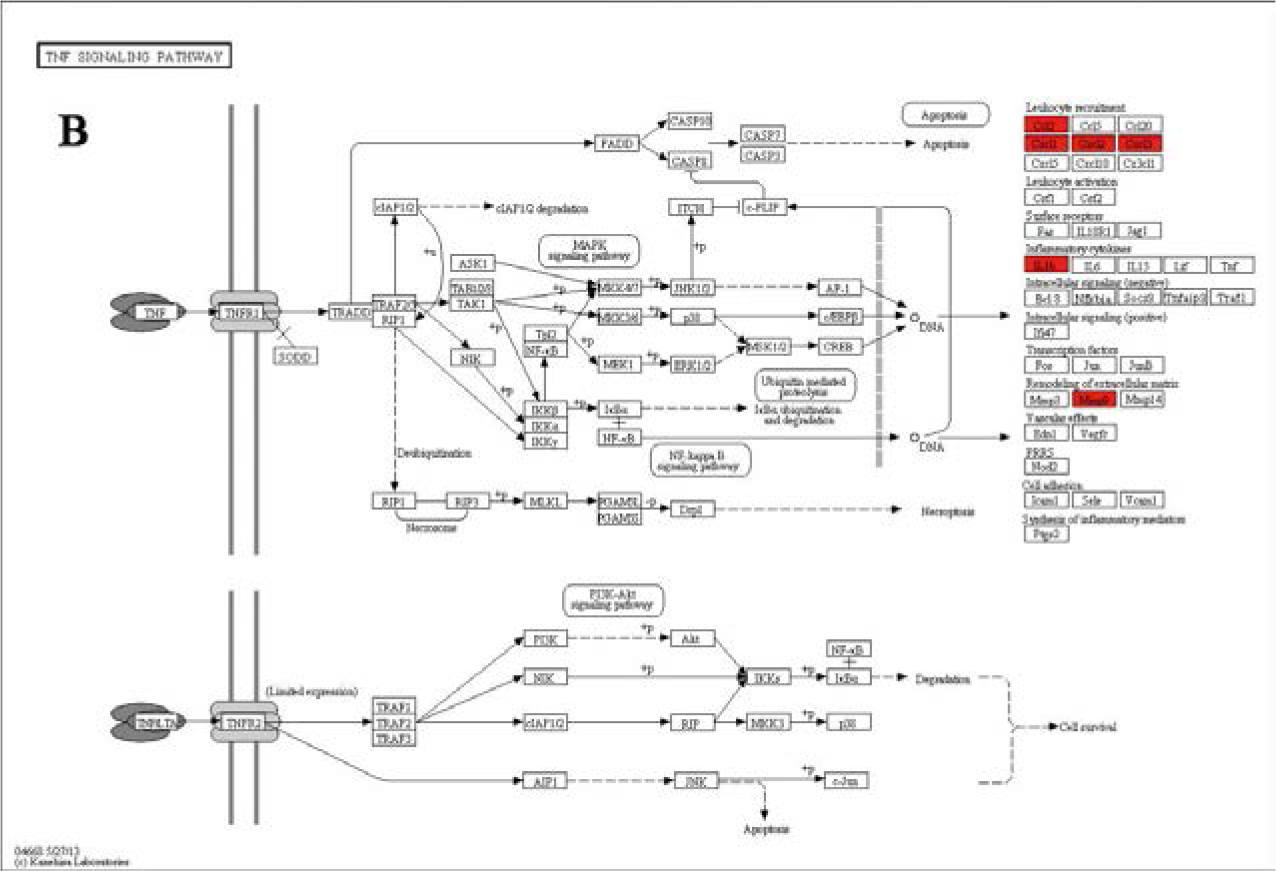
**A:** chemokine signaling pathway. **B:** tumor necrosis factor signaling pathway. All signaling pathways are cited from www.kegg.jp/kegg/kegg1.html.

### Expression Analysis by qRT-PCR

Based on the RNA deep sequencing analysis of the expression, five genes, including CXCL1, CXCL2, CXCL3, CCL7 and MMP9, were selected for confirmation as well as to monitor their expression with qRT-PCR, and the data was statistically analyzed as follows (figure 9).

**Figure 9.**
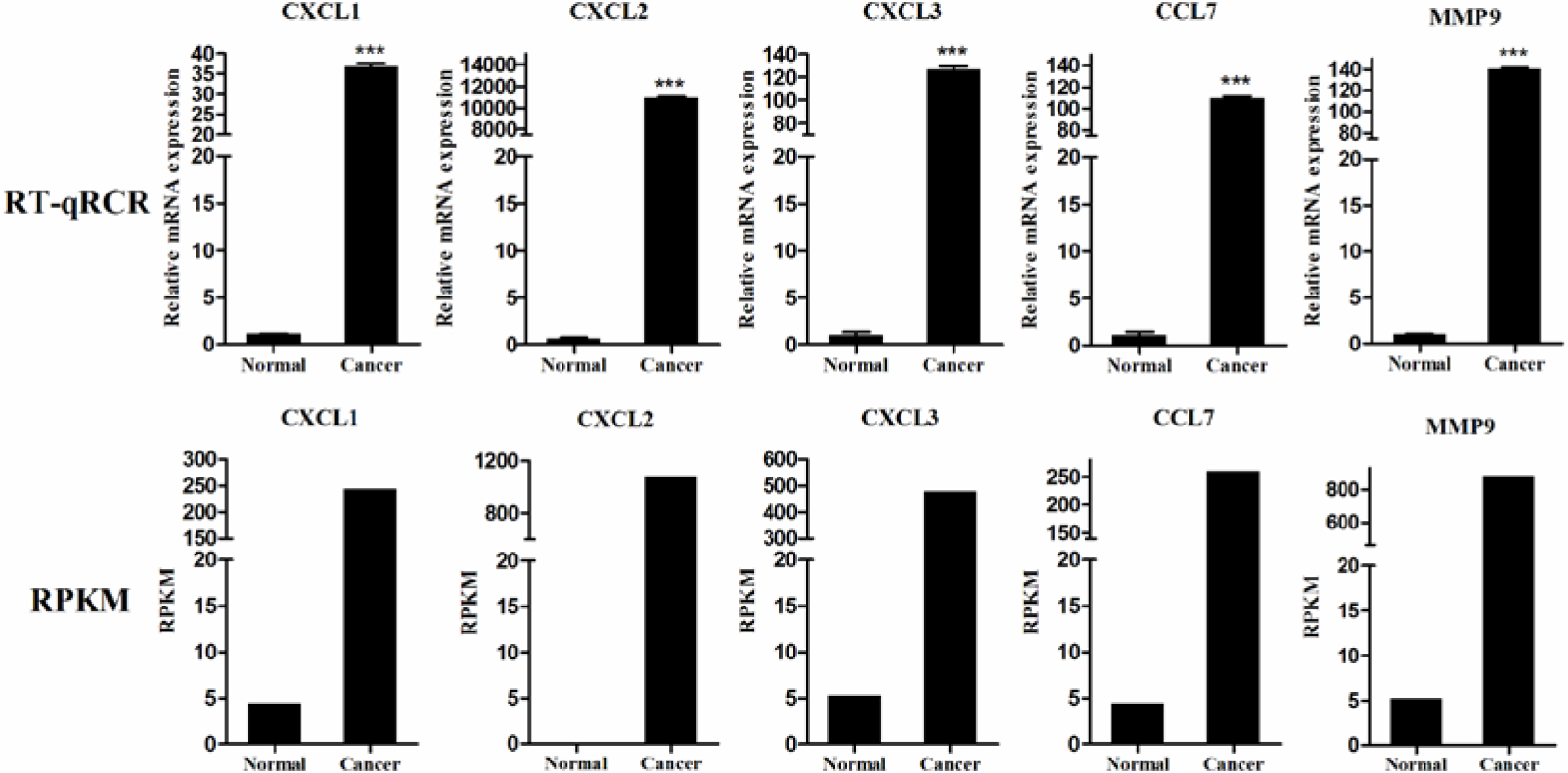
qRT-PCR validation transcriptome sequencing results. The first line is the relative expression level of differentially expressed genes in oral buccal mucosa tissues of Chinese hamster detected by qRT-PCR (n=3).The second line is the expression level of differentially expressed genes in each tissue detected by transcriptome sequencing (RPKM, n=3).

Compared with normal group, in the cancer group the mRNA expression level of CXCL1, CXCL2, CXCL3, CCL7 and MMP9 increased remarkably. Meanwhile the transcriptome sequencing revealed that the differentially expressed genes: CXCL1, CXCL2, CXCL3, CCL7 and MMP9 were also significantly up-regulated in the cancer group compared to the normal group. The relative expression of mRNA detected by qRT-PCR and the transcriptome sequencing data (RPKM) were highly consistent, indicating that the sequencing data was reliable, suggesting that CXCL1, CXCL2, CXCL3, CCL7 and MMP9 may promote the development, invasion and metastasis of OSCC.

## Discussion

Oral cancer is the most common malignancy of head and neck cancer in the world and it is a leading cause of cancer-related mortality. Epidemiological data have shown that high risk factors, such as tobacco, alcohol and Human papillomavirus infection are closely related to OSCC (Chi et al., 2015). Recently, the incidence of oral cancer is rising with ages every year and more seriously, the patients are mostly diagnosed at a relatively late stage which lead to increase in the cost of treatment and decrease in treatment outcomes(Oh et al.). In the present biological era, the prognosis of this deadly disease has improved to some extent because of the enhancement of technologies (Frédérique et al., 2012), but diagnosing the tumor at its initial stages relapse, and metastasis are the major challenge to improve the scope of patient survival (Dahiya and Dhankhar, 2016). Therefore, it is urgent and imperative to delve into the molecular mechanisms of the development and progression of oral cancer to guide clinical treatment and prognosis.

Since Sally first established the DMBA-induced oral carcinogenesis model in the cheek pouch of hamster in 1954, it has become a classic animal model of OSCC (Salley, 1954). In present study, we successfully constructed the animal model of OSCC using tri-weekly applications of a 5% solution of DMBA in acetone onto the cheek pouch of Chinese hamsters over about a 15-week period. For models like the hamster model for OSCC, high-throughput sequencing provides a powerful tool for analyzing both mRNA expression patterns and quantitative expression levels, as it profiles thousands of genes simultaneously. New high-throughput sequencing technologies have enabled the detection of novel transcripts through increased sensitivity. These recent advances have facilitated more comprehensive and more thorough research into the effects of transcription and translation (Zeng JH et al., 2019). This technology is much more efficient than the now outmoded and time-consuming methods used in earlier work, and is becoming the broadest transcriptome research tool available.

Different from earlier studies, our study considered the development and progression of OSCC holistically, including a variety of pathways and genes, rather than just a single factor. Using high-throughput sequencing analysis, we evaluated mRNA expression profiles of OSCC and normal cheek pouch mucosa tissues, and identified 128 mRNAs that were up-regulated and 66 down-regulated in cancer tissues compared with normal tissues. What’s more,because genes often cooperate with each other to perform their biological functions, pathway analyze facilitate further understanding genes and their roles. By the KEGG enrichment analysis, we found that differentially expressed genes were enriched in 317 pathways, of which 8 pathways were significantly enriched. In addition, we used GO analysis to screen genes which were significantly enriched in pathways, and selected five genes(*CXCL1*, *CXCL2*, *CXCL3*, *CCL7*, and *MMP9*) that were highly expressed in OSCC, may be closely related to the development of oral cancer and significantly enriched in chemokine signaling pathway and TNF signaling pathway for qRT-PCR to validate transcriptome sequencing. The GO functional enrichment analysis showed that the differentially expressed mRNAs in OSCC were mainly enriched in cellular components, including extracellular matrix, extracellular space and extracellular region, etc; involved molecular functions that included cytokine activity and extracellular matrix structural constituent; involved biological processes that included granulocyte chemotaxis, granulocyte migration, cell chemotaxis, cell migration, cell motility and localization of cell, etc. RT-qPCR confirmed that the expression of *CXCL1*, *CXCL2*, *CXCL3*, *CCL7* and *MMP9* were consistent with RNA-seq. Meanwhile, qRT-PCR confirmed that *CXCL2* had the largest differential expression folds in OSCC tissues, revealing that *CXCL2* may play a central role in the molecular mechanism of OSCC. The high expression of *CXCL2* suggests that it can be used as a new target for diagnosis, treatment and prognosis of OSCC, and it also provides novel ideas and important theoretical basis for the search for potential molecular markers of oral cancer. These findings suggest that there is still an inflammatory response during OSCC and the previous differential expression analysis also reflected that chemokine is still highly expressed in OSCC. In summary, the inflammatory response is in an uncontrollable state and it is difficult to restore the body’s homeostasis, eventually resulting in the production of tumors.

Chemokines are a group of small proteins(8–12 KDs), responsible for transmitting signals for cell migration, inflammation regulation, and angiogenesis, classifed into four subgroups referred to as CXC/α, CC/β, CX3C/δ or C/δ families (Baggiolini, 1998; Zlotnik and Yoshie, 2000). They are mostly known for their role in immunesurveillance and infammatory responses, but they have been also implicated in many pathological processes of malignant tumor (Mahboobeh et al., 2014). Recently, the research of chemokines in the oncology has drawn extensive attention. Meanwhile, related studies have demonstrated the involvement of chemokines and their receptors in various tumors such as liver cancer, nasopharyngeal cancer, and breast cancer (Jing et al., 2016; Shah et al., 2015; Weitzenfeld et al., 2016). Abnormal function of chemokines in cancer promotes cell survival, facilitated proliferation, angiogenesis, and metastasis in multiple types of tumors (Paola et al., 2011). The research of Vera Levina indicate that chemokines and growth factors produced by tumors by binding to the cognate receptors on tumor and stroma cells could provide proliferative and anti-apoptotic signals helping tumors to escape drug-mediated destruction (Vera et al., 2010). Furthermore, it is believed that chronic inflammatory conditions facilitate oral carcinogenesis, and functions of cytokine-dependent and chemokine-dependent immunoregulatory pathways are apparent in oral carcinoma.

Among them, Jung(Da-Woon et al., 2010) confirmed that *CCL7* may participate in the invasion and metastasis of OSCC through a molecular dialogue with the *CCR1* and *CCR3*. In this study, the chemokine *CXCL2*, *CXCL3*, and *CXCL1* were all highly expressed in OSCC, and Peng (Peng et al., 2015)also detected significant upregulation of *CXCL2* and *CXCL3* by microarray analysis of OSCC in rats, indicating abnormal expression of *CXCL1*, *CXCL2* and *CXCL3* is an important factor leading to pathological changes in the oral cancer model. Furthermore, OSCC microarray analysis also reflected abnormal expression of *CXCL1*, *CXCL2* and *CXCL3* in tumor tissues (Sanjukta et al., 2015), while clinical and follow-up basic studies on *CXCL1*, *CXCL2*, and *CXCL3* are rare in oral cancer and most of them are found in liver cancer, prostate cancer and breast cancer (Gui et al., 2016; Li et al., 2015; See et al., 2014). Thence, the further study of these significantly different chemokines in the pathogenesis has important implications for the diagnosis and treatment of oral cancer.

Matrixmetallo-proteinases (MMPs) are major proteolytic enzymes that are involved in the normal extracellular matrix (ECM) turnover. Its main function is to degrade and remodel the ECM, maintain the dynamic balance of ECM, and participate in many pathological and physiological processes in the body. Many of clinical studies have confirmed the high expression of *MMP9* and *MMP13* in OSCC (Jingqiu et al., 2015; Monteiro et al., 2016; Nanda et al., 2014), which is consistent with our research. Monteiro (Monteiro et al., 2016) applied immunohistochemistry to detect the expression of MMP9 protein in 60 cases of OSCC and found that *MMP9* was strongly expressed in cytoplasm of tumor cell in 83.7% of patients, suggesting that *MMP9* is a potential tumor marker of oral cancer. Yu (Yu et al., 2011) knocked out the chemokine receptor *CXCR4* gene in Tca8113 cells, as a result, the expression levels of *MMP9* and *MMP13* were significantly reduced, and the invasion and metastasis of cells were weakened, suggesting that *MMP9* and *MMP13* play a pivotal role in the invasion and metastasis of OSCC. The matrix metalloproteinase we screened is *MMP9*, which is enriched with *CXCL1*, *CXCL2*, and *CXCL3* in TNF signaling pathway, suggesting that *MMP9* may promote tumorigenesis through the interaction with chemokines. Gao et al. (Gao et al., 2015) said that *CXCL5*/*CXCR2* axis activates the PI3K/AKT signaling pathway of the tumor signaling pathway, leading to the up-regulation of *MMP2* and *MMP9*, which promotes the metastasis of bladder cancer, confirming the interaction between *MMP9* and chemokines. However, the specific interaction between *MMP9* and *CXCL1*, *CXCL2*, *CXCL3* in OSCC still needs further study. Judging from these figures, we can draw the conclusion that the expression trend of the five genes in qRT-PCR was consistent with the transcriptome sequencing, which validated the reliability of RNA-seq data and further demonstrated that the chemokine signaling pathway and TNF signaling pathway may closely related to the occurrence and development of OSCC.

To our knowledge, our study presents the first genomewide profiling of mRNAs of squamous cell carcinoma in oral buccal pouch of Chinese hamster by high-throughput RNA-Seq. In addition, our findings support previous studies reporting that the progression of OSCC was influenced by chemokines, suggesting that chemokine signaling pathway, Cytokine-cytokine receptor interaction signaling pathway, and TNF signaling pathway may play a central role in the invasion and metastasis of OSCC (Farkas et al., 2013; Toung et al., 2011). Peeping a spot to see overall picture: local delicate change was packed with the complex issues of the whole organism. But for the further verification and exploration, researches on the cellular level and the significant expression of proteins during the development and progression of OSCC should be carried out.

## Conclusions

All the findings, including the chemokine signaling pathway, TNF signaling pathway and classic genes provided an extensive bioinformatics analysis of DEGs and revealed a series of targets and pathways, which may affect the carcinogenesis and progression of OSCC, for future investigation. These findings add to significant insights into the diagnosis and treatment of this disease. Meanwhile, it should be noted that this study examined the DEGs in oral cancer Chinese hamster animal model by qRT-PCR and bioinformatics analysis; further research needs to be done to explore more specific mechanisms. Notwithstanding its limitation, these findings significantly improved the understanding of underlying molecular mechanisms in OSCC, and the key genes and pathways could be used as diagnostic and therapeutic targets and diagnostic biomarkers.

## Acknowledgements

Thank University of Shanxi Medical University for offering a powerful bioinformatics platform for our study. Thank Professor Guohua Song for her careful guidance.

## Author contributions

G.H.S. designed the study and contributed funding. G.Q.X., J.N.W., and B.HF. established animal model, completed RNA sequencing and statistical analyze. L.F.X, X.T.W., J.P.G. AND R.J.X. collected samples and processed samples. Z.Y.C. provided the necessary tools and instruments for the experiments. G.Q.X. and J.P.G. contributed to writing the manuscript. All authors discussed the results and commented on the manuscript.

## Reference

Baggiolini, M. (1998). Chemokines and leukocyte traffic. Nature 392, 565–568.

Chi, A. C., Day, T. A. and Neville, B. W. (2015). Oral cavity and oropharyngeal squamous cell carcinoma--an update. Ca A Cancer Journal for Clinicians 65, 401–421.

Christopher, G., Jiangwen, Z., Butler, J. E., David, H. and Catherine, D. (2010). Sex-specific parent-of-origin allelic expression in the mouse brain. Science 329, 682–685.

Da-Woon, J., Min, C. Z., Jinmi, K., Kyungshin, K., Ki-Yeol, K., Darren, W. and Jin, K. (2010). Tumor-stromal crosstalk in invasion of oral squamous cell carcinoma: a pivotal role of CCL7. International Journal of Cancer 127, 332–344.

Dahiya, K. and Dhankhar, R. (2016). Updated overview of current biomarkers in head and neck carcinoma. World Journal of Methodology 6, 77–86.

Farkas, M. H., Grant, G. R., White, J. A., Sousa, M. E., Consugar, M. B. and Pierce, E. A. (2013). Transcriptome analyses of the human retina identify unprecedented transcript diversity and 3.5 Mb of novel transcribed sequence via significant alternative splicing and novel genes. Bmc Genomics 14, 486–486.

Ferlay, J., Soerjomataram, I., Dikshit, R., Eser, S., Mathers, C., Rebelo, M., Parkin, D. M., Forman, D. and Bray, F. (2015). Cancer incidence and mortality worldwide: sources, methods and major patterns in GLOBOCAN 2012. International Journal of Cancer 136, E359–E386.

Frédérique, N., Jean-Charles, S. and Fabien, C. (2012). Tumour molecular profiling for deciding therapy-the French initiative. Nature Reviews Clinical Oncology 9, 479–486.

Gao, Y., Guan, Z., Chen, J., Xie, H., Yang, Z., Fan, J., Wang, X. and Li, L. (2015). CXCL5/CXCR2 axis promotes bladder cancer cell migration and invasion by activating PI3K/AKT-induced upregulation of MMP2/MMP9. International Journal of Oncology 47, 690–700.

Gibb, E. A., Enfield, K. S., Tsui, I. F., Chari, R., Lam, S., Alvarez, C. E. and Lam, W. L. (2010). Deciphering squamous cell carcinoma using multidimensional genomic approaches. Journal of Skin Cancer 2011, 541405.

Gimenez-Conti IB and Tj, S. (1993). The hamster cheek pouch carcinogenesis model. J Cell Biochem 1, 83–90.

Gui, S. L., Teng, L. C., Wang, S. Q., Liu, S., Lin, Y. L., Zhao, X. L., Liu, L., Sui, H. Y., Yang, Y. and Liang, L. C. (2016). Overexpression of CXCL3 can enhance the oncogenic potential of prostate cancer. International Urology & Nephrology 48, 1–9.

Jing, Y., Xing, L., Chen, J., Xie, C., Xia, W., Chen, J., Zeng, T., Ye, Y., Ke, L. and Yu, Y. (2016). CCL2-CCR2 axis promotes metastasis of nasopharyngeal carcinoma by activating ERK1/2-MMP2/9 pathway. Oncotarget 7, 15632–15647.

Jingqiu, B., Xi, B., Bing, L., Fei, C. and Peng, C. (2015). Inhibition of metastasis of oral squamous cell carcinoma by anti-PLGF treatment. Tumour Biology the Journal of the International Society for Oncodevelopmental Biology & Medicine 36, 2695.

Kanehisa, M., Furumichi, M., Mao, T., Sato, Y. and Morishima, K. (2017). KEGG: new perspectives on genomes, pathways, diseases and drugs. Nucleic Acids Research 45, D353–D361.

Kang, M. K. and Park, N. H. (2001). Conversion of normal to malignant phenotype: telomere shortening, telomerase activation, and genomic instability during immortalization of human oral keratinocytes. Crit Rev Oral Biol Med 12, 38–54.

Lee, S.-H., Singh, I., Tisdale, S., Abdel-Wahab, O., Leslie, C. S. and Mayr, C. (2018). Widespread intronic polyadenylation inactivates tumour suppressor genes in leukaemia. Nature 561, 127–131.

Li, L., Xu, L., Yan, J., Zhen, Z. J., Ji, Y., Liu, C. Q., Wan, Y. L., Zheng, L. and Xu, J. (2015). CXCR2–CXCL1 axis is correlated with neutrophil infiltration and predicts a poor prognosis in hepatocellular carcinoma. Journal of Experimental & Clinical Cancer Research 34, 1–10.

Li, X., Chen, J., Hu, X., Huang, Y., Li, Z., Zhou, L., Tian, Z., Ma, H., Wu, Z., Chen, M. et al. (2011). Comparative mRNA and microRNA expression profiling of three genitourinary cancers reveals common hallmarks and cancer-specific molecular events. Plos One 6, e22570–e22570.

Lodi, G., ., Franchini, R., ., Bez, C., ., Sardella, A., ., Moneghini, L., ., Pellegrini, C., ., Bosari, S., ., Manfredi, M., ., Vescovi, P., . and Carrassi, A., . (2010). Detection of survivin mRNA in healthy oral mucosa, oral leucoplakia and oral cancer. Oral Diseases 16, 61–67.

Mahboobeh, R., Fahimeh, A., Mousa, T., Ali, M., Nooshafarin, C. and Abbas, G. (2014). Expression of chemokines and chemokine receptors in brain tumor tissue derived cells. Asian Pacific Journal of Cancer Prevention Apjcp 15, 7201–5.

Monteiro, L., Delgado, M., Ricardo, S., Amaral, B., Salazar, F., Pacheco, J., Lopes, C., Bousbaa, H. and Warnakulasuryia, S. (2016). Prognostic significance of CD44v6, p63, podoplanin and MMP‐9 in oral squamous cell carcinomas. Oral Diseases 22, 303–312.

Nanda, D. P., Dutta, K., ., Ganguly, K. K., Hajra, S., ., Mandal, S. S., Biswas, J., . and Sinha, D., . (2014). MMP-9 as a?potential biomarker for carcinoma of oral cavity: a?study in?eastern India. Neoplasma 61, 747–757.

None. (1978). Definition of leukoplakia and related lesions: An aid to studies on oral precancer. Oral Surgery Oral Medicine & Oral Pathology 46, 518,539–537,539.

Oh, H. N., Oh, K. B., Lee, M. H., Seo, J. H., Kim, E., Yoon, G., Cho, S. S., Cho, Y. S., Choi, H. W. and Chae, J. I. JAK2 regulation by Licochalcone H inhibits the cell growth and induces apoptosis in oral squamous cell carcinoma. Phytomedicine.

Paola, A., Giovanni, G., Federica, M. and Alberto, M. (2011). Chemokines in cancer related inflammation. Experimental Cell Research 317, 664–673.

Peng, X., Li, W., Johnson, W. D., Torres, K. E. and Mccormick, D. L. (2015). Overexpression of lipocalins and pro-inflammatory chemokines and altered methylation of PTGS2 and APC2 in oral squamous cell carcinomas induced in rats by 4-nitroquinoline-1-oxide. Plos One 10, e0116285.

Raimondi, A., Cabrini, R. and Itoiz, M. E. (2005). Ploidy analysis of field cancerization and cancer development in the hamster cheek pouch carcinogenesis model. Journal of Oral Pathology & Medicine 34, 227–231.

Rajasekar, M., Suresh, K. and Sivakumar, K. (2016). Diosmin induce apoptosis through modulation of STAT-3 signaling in 7,12 dimethylbenz(a)anthracene induced harmster buccal pouch carcinogenesis. Biomedicine & Pharmacotherapy 83, 1064–1070.

Ramu, A., Kathiresan, S. and Bakrudeen, A. (2017). Gramine inhibits angiogenesis and induces apoptosis via modulation of TGF-β signaling in 7,12 dimethylbenz[a]anthracene (DMBA) induced hamster buccal pouch carcinoma. Phytomedicine International Journal of Phytotherapy & Phytopharmacology 33, 69.

Salley, J. J. (1954). Experimental carcinogenesis in the cheek pouch of the Syrian hamster. Journal of Dental Research 33, 253.

Sanjukta, C., Shaleen, M., Jyoti, D. and Dhananjaya, S. (2015). Whole genome expression profiling in chewing-tobacco-associated oral cancers: a pilot study. Medical Oncology 32, 60.

See, A. L., Chong, P. K., Lu, S. Y. and Lim, Y. P. (2014). CXCL3 is a potential target for breast cancer metastasis. Current Cancer Drug Targets 14, -.

Shah, A. D., Bouchard, M. J. and Shieh, A. C. (2015). Interstitial Fluid Flow Increases Hepatocellular Carcinoma Cell Invasion through CXCR4/CXCL12 and MEK/ERK Signaling. Plos One 10, e0142337.

Shlok, G., Weidong, K., Yingwei, P., Qun, M. and Mackillop, W. J. (2010). Temporal trends in the incidence and survival of cancers of the upper aerodigestive tract in Ontario and the United States. International Journal of Cancer 125, 2159–2165.

Siegel, R., Ma, J., Zou, Z. and Jemal, A. (2014). Cancer statistics, 2014. Ca Cancer J Clin 64, 9.

Toung, J. M., Morley, M., Li, M. and Cheung, V. G. (2011). RNA-sequence analysis of human B-cells. Genome Research 21, 991.

Vera, L., Yunyun, S., Brian, N., Xiaoning, L., Yuri, G., Melissa, L., Anna, L. and Elieser, G. (2010). Chemotherapeutic drugs and human tumor cells cytokine network. International Journal of Cancer 123, 2031–2040.

Weitzenfeld, P., Kossover, O., Körner, C., Meshel, T., Wiemann, S., Seliktar, D., Legler, D. F. and Ben-Baruch, A. (2016). Chemokine axes in breast cancer: factors of the tumor microenvironment reshape the CCR7-driven metastatic spread of luminal-A breast tumors. Journal of Leukocyte Biology 99, 1009–1025.

Yu, T., Wu, Y., Helman, J. I., Wen, Y., Wang, C. and Li, L. (2011). CXCR4 promotes oral squamous cell carcinoma migration and invasion through inducing expression of MMP-9 and MMP-13 via the ERK signaling pathway. Molecular Cancer Research Mcr 9, 161.

Zeng JH, Lu W, Liang L, Chen G, Lan HH, Liang XY and X, Z. (2019). Prognosis of clear cell renal cell carcinoma (ccRCC) based on a six-lncRNA-based risk score: an investigation based on RNA-sequencing data. J Transl Med 17, 281.

Zhu, J., Jiang, Z., Gao, F., Hu, X., Zhou, L., Chen, J., Luo, H., Sun, J., Wu, S. and Han, Y. (2011). A systematic analysis on DNA methylation and the expression of both mRNA and microRNA in bladder cancer. Plos One 6, e28223.

Zlotnik, A. and Yoshie, O. (2000). Chemokines : A New Classification System and Their Role in Immunity. Immunity 12, 121–127.

